# A model of sediment retention by vegetation for Great Britain: new methodologies & validation

**DOI:** 10.1101/2023.08.17.553678

**Authors:** Danny A.P. Hooftman, James M. Bullock, Paul M. Evans, John W. Redhead, Lucy E. Ridding, Varun Varma, Richard F. Pywell

**Affiliations:** Lactuca, Environmental Data Analyses and Modelling, 1112NC, Diemen, The Netherlands; UK Centre for Ecology & Hydrology, OX10 8BB, Wallingford, UK; Rothamsted Research, AL5 2JQ, Harpenden, UK

**Keywords:** Ecosystem Services, Erosion, Erosivity, InVEST, Soil Retention, Sediments, Vegetation Cover, United Kingdom

## Abstract

Soil erosion is an substantial environmental concern worldwide. It has been historically and is of increasingly concern currently. Next to natural processes, over 2 million hectares of soil are at risk of erosion through intensifying agriculture in the Great Britain (England, Wales, Scotland and their territorial islands). Predictive soil erosion models, in the form of Ecosystem Service tools, aid in helping to identify areas that are vulnerable to soil erosion. Yet, no predictions for erosion or sediment retention by vegetation based on local data have been developed for Great Britain or the United Kingdom as a whole.

Here we develop an erosion retention model using the InVEST platform, which is based on the RUSLE mathematical framework. We parameterise the model, as far as feasible, with GB specific input data. The developed model estimations are validated against suspended solids concentrations (sediments) in throughout England and Wales.

Next to presenting the first GB wide estimate of erosion and erosion retention using the InVEST SDR module, we test three approaches here that differ from more widely applicable RUSLE model inputs, such as created for Europe as a whole. Here, we incorporate (1) periodicity to allow erosion to potentially fluctuate within years; (2) GB-specific cover periodic management factors estimates, including a range of crop types, based on observed satellite NDVI values (3) soil erosivity under heavy rainfall following GB estimates for 2000-2019.

We conclude that both the GB created erosivity layer as the added periodicity do not seem to be provide substantial improvement over non-periodic estimated created with more widely available data, when validated against this set of suspended solids in rivers. In contrast, the observed cover management factors calculated from NDVI are a good improvement affecting the ranking order among catchments. Therefore, the generating of cover management factors using NDVI data could be promoted as method for InVEST SDR model development and in more general for developing RUSLE-based erosion estimates worldwide.

## 1. Introduction

Soil erosion has been an environmental concern for millennia (Morgan 2005). Healthy soil is the foundation of agriculture and an essential resource to ensure human needs throughout time (Haygarth & Ritz 2009), even more in the intensifying 21st century (Borrelli *et al*. 2017). Although a natural process, soil erosion is aggravated by anthropogenic activities (Adornado *et al*. 2009; Benavidez *et al*. 2018). As a consequence, soil erosion by water has become one of the major threats to soils worldwide (FAO 2015) in Europe (Panagos *et al*. 2015a; 2021) and the United Kingdom (Haygarth & Ritz 2009; Boardman *et al*. 2017). Impacts of soil erosion include negative effects on delivery of many natural processes such as crop production, water storage and filtration, nutrient cycling, biodiversity and carbon storage (Panagos *et al*. 2015a; Robinson *et al*. 2017; Rust *et al*. 2022).

For agricultural areas, not only does it wash away potential crop or grazing areas, but through removal of the topsoil many of the supporting nutrients for following years are permanently lost, being washed with the streams. The European Union has listed soil erosion among the key soil threats listed within the Soil Thematic Strategy of the European Commission (EC 2012; Montanarella 2011; Panagos *et al*. 2015a) with soil loss by water erosion projected to increase by 20% in EU and UK by 2050 (Panagos *et al*. 2021). In the UK, over 2 million hectares of soil are at risk of erosion while intensive agriculture has caused arable soils to lose about 40 to 60% of their organic carbon (EA 2023). Where Europe wide predictive estimations have been published (Panagos *et al*. 2015a, 2021), as yet no recent UK-local data driven sediment retention or erosion predictions have been developed for the full UK or Great Britain –the UK without Northern Ireland–..

Predictive soil erosion models aid in helping to identify areas that are vulnerable to soil erosion, estimated potential erosion rates, and highlight causes of soil erosion. As well as aid in identifying They range from relatively simple empirical models, conceptual models, to complex physics-mathematical models (Benavidez *et al*. 2018). Karydas *et al*. (2014) identified 82 water-erosion models over different spatial/temporal scales and with various levels of complexity (Panagos *et al*. 2015a), a number that has been growing over the last years (Borrelli *et al*. 2021), also due to the increasing availability of Ecosystem Services model platforms (Willcock *et al*. 2023). The most commonly used complex mathematical framework to predict erosion is the Universal Soil Loss Equation (USLE; Wischmeier and Smith, 1978) and its revised form (RUSLE; Renard *et al*., 1997). (R)USLE estimates long-term average for annual soil loss by rainfall and overland erosion (Panagos *et al*. 2015a). The approach has been used for many areas and is the dominant approach used worldwide (Panagos *et al*. 2014; Benavidez *et al*. 2018; Kumar *et al*. 2022)

One of the modelling platform that developed a RUSLE based approach is InVEST (NatCap 2023), the world’s leading platform for Ecosystem Service estimations since 2009 (Ochoa & Urbina-Cardona 2017; Bratman *et al*. 2020). This RUSLE-based InVEST Sediment Delivery Ratio (SDR) model is an spatially explicit erosion model focussing on overland erosion and soil retention by vegetation, i.e., avoided erosion. Outputs from the model include the sediment load delivered to the stream at an annual time scale, as well as the amount of sediment retained by vegetation and topographic features.

However, to our knowledge an recent InVEST SDR model assessment focussed on the UK, or parts therefore, has not been published. In this paper we parametrise an InVEST model, driven by local data, for Great Britain (England, Wales, Scotland and their territorial islands) and present as such full GB assessment for the first time of amount of estimated erosion. More importantly, we highlight the areas where retention capacity by vegetation is limited. The developed model estimations will be validated against suspended solids concentrations (sediments) in throughout England and Wales.

Moreover, we will use our model approach to test for three data input approaches, hence our paper has a strong methodological focus. InVEST requires bespoke input data for all data-sets, i.e., it has no framework driven ‘automatic’ data inputs as ARIES or Co$ting Nature have, as large Ecosystem Service platforms (Willcock *et al*. 2023; Bagstad *et al*. 2013). However, how elaborate such inputs are, is up to the user. Secondly, InVEST is and annual model, where crop cover varies over the year, which influences the amount of soil erosion through temporally changing cover retention when rainfall is dyssynchronous throughout the year. Here we focus on generating highly GB specific inputs and use the model such that such temporally variable cover can be included. We test the level of validation accuracy of those input approaches against a best-estimate base-line approach following inputs derived from literature, which are generally not locally specific.

Next to presenting the first GB wide SDR model driven by local data, we are testing the following new approaches, they will be explained more below:

1. Incorporating periodicity to allow erosion to potentially fluctuate within years, due to interacting temporally larger erosivity and lowered soil coverage by vegetation (i.e., in winter);
2. GB-specific cover management C-factors estimates (both periodic and non-periodic), including a range of crop types, based on observed satellite NDVI values as proxy for soil coverages;
3. Erosivity (R; periodic and non-periodic) following GB estimates for 2000-2019 in combination with a published regression approach.

## 2. Methods and new elements

All calculation were done using InVEST 3.12.0, ArcGIS Pro 3.0.3 and Matlab v7.14.0.739. Codes can be found on https://github.com/dhooftman72/AgLandSDR. The area of study is Great Britain (GB), defined as the United Kingdom (UK) without Northern Ireland and the Isle of Man, but including the outer islands such as the Hebrides, Shetlands, and Orkneys (Figure 1a).

**Figure 1.**
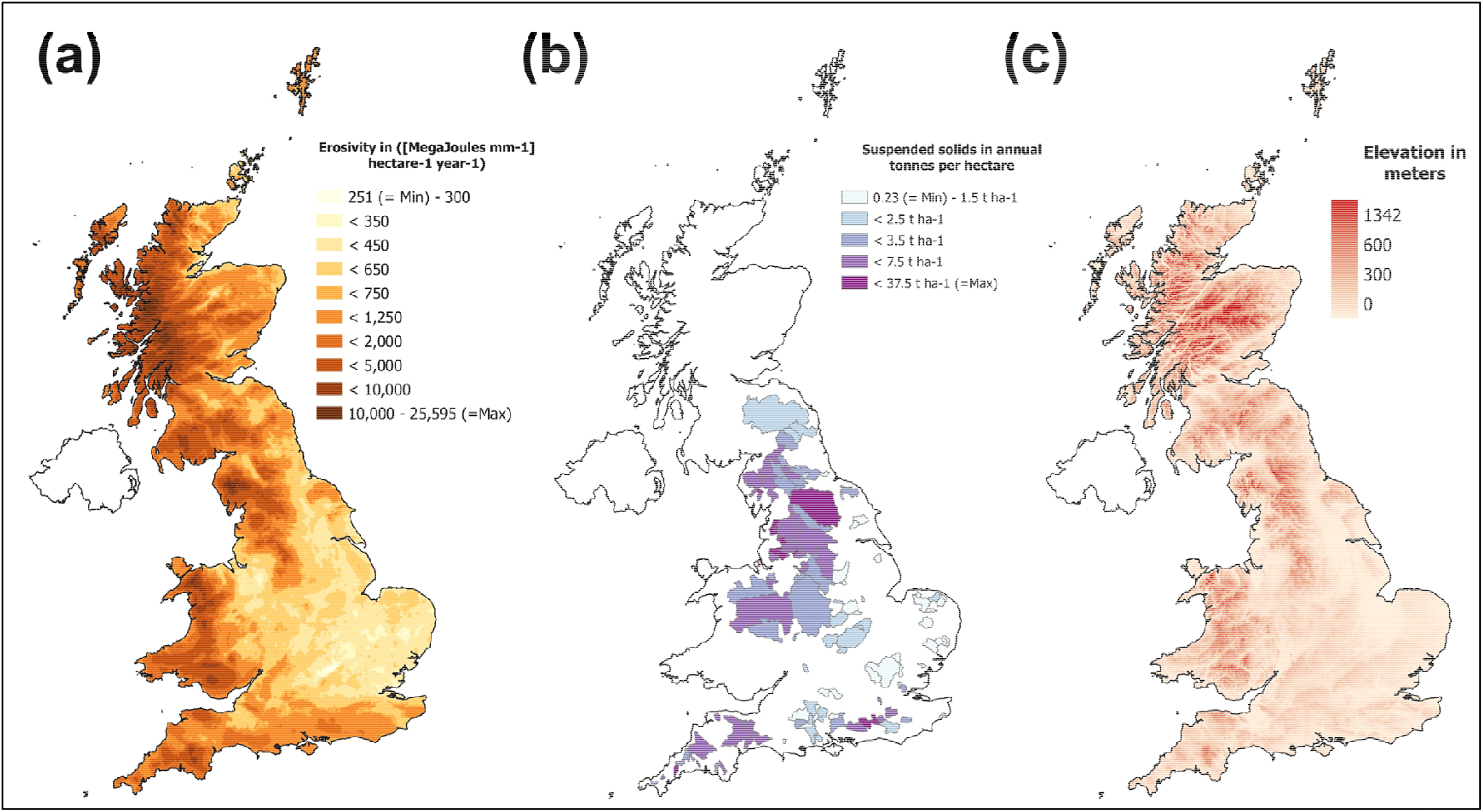
Methods data. (A) Calculated annual erosivity layer for GB from daily CEH-GEAR data (Tanguy *et al*. 2021) using the regression approach from Ferreira & Panagopoulos (2014). (B) The 178 catchments used for validation– with a matching outlet locations for both multi-annual water quality monitoring and NFRA water discharge measurements. Provided is the annual amount of suspended solids (“sediments”) from the UK water quality archive (“Wims”). Catchments are delineated to monitoring locations and can partly overlap. (C) Elevation following Moore & Flavin (1990; 1994).

### 2.1 The InVEST Erosion Model

The InVEST SDR model is an erosion model focussing on overland erosion, it does not model gully, bank, or mass erosion (NatCap 2023). Outputs from the model include the sediment load delivered to the stream at an annual time scale, as well as the amount of sediment eroded in the catchment and retained by vegetation and topographic features. The SDR model is a spatially explicit model working at the spatial resolution of the inputted digital elevation model (DEM) raster. For each pixel, the model first computes the amount of annual soil loss from that pixel (Eq. 1), then computes the proportion of soil loss actually reaching the stream (Eq. 6). Once sediment reaches the stream, it is assumed to be delivered to the catchment outlet; thus no in-stream processes which could increase or decrease sediment loads are modelled. This approach was proposed by Borselli *et al*. (2008) and has had much following (InVEST calculations are reported in *e.g*., Cavalli *et al*. 2013; Ougougdal *et al*. 2020; Gashaw *et al*. 2021; Marquez *et al*. 2021). However, to our knowledge an InVEST SDR model assessment for the UK, or parts therefore, has not been published.

#### 2.1.1 Annual soil loss per pixel

The amount of annual soil loss per pixel (*A*_*i*_) in tons per hectare per year (t h^-1^ y^-1^) and is calculated by the Revised Universal Soil Loss Equation (RUSLE; Renard *et al*. 1997; Panagos *et al*. 2015a; Benavidez *et al*. 2018), with is a linear correction model calculating the potential run-off (*R, K, L* and *S* factors) limited by vegetation and management actions (C & P factors). In equation for gridcell *i* this follows:

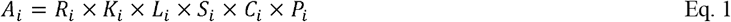

with:

*A*_*i*_ is mean annual soil loss (tons sediment hectare^-1^ year^-1^);

*R*_*i*_ the rainfall erosivity ([MegaJoules mm^-1^] hectare^-1^ hour^-1^ year^-1^; see below);

*K*_*i*_ the soil erodibility (metric ton hours per [MJ mm^-1^]; see below);

*L*_*i*_ is the slope length factor (unitless; see NatCap 2003; Benavidez *et al*. 2018);

*S*_*i*_ is the slope steepness factor (unitless; see NatCap 2003; Benavidez *et al*. 2018);

*C*_*i*_ is the cover management factor (unitless; see below);

*P*_*i*_ is the support practice factor (unitless; see below).

Within InVEST *L*_*i*_ and *S*_*i*_ are calculated based on a provided DEM. In contrast, *R*_*i*_, *K*_*i*_ and need to be provided in mapped form, whereas *C*_*i*_ and *P*_*i*_ need to be provided in tabled form for predefined land cover classes. Therefore, a Land Cover Map needs to be provided to InVEST, matching the table.

#### 2.1.2. Periodic soil loss per pixel

Although *A*_*i*_ following RUSSLE, is an annual number, we suggest here it could be split into periods as in needed below to incorporate cover-management factors (*C*_*i*_) that vary throughout the year. This is following the reasoning that the *R*_*i*_ factor is the sum of all erosive events (*j*) throughout the year exceeding a threshold of rainfall intensity and length (Renard *et al*. 1997; Benavidez *et al*. 2018).

Therefore Eq. 1 could be rewritten as sum of all erosive events for gridcell *j* with corresponding time-specified cover management factor (*C*_*ij*_), *i.e*., the vegetation cover at the moment of rainfall as:

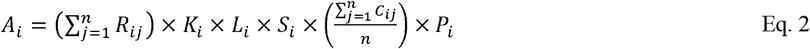

With parameters as Eq. 1, *j* an erosive event and n the total of erosive events

When extending this reasoning, 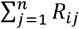could denote a sum of individual erosive events but also a ĵsum of periods (in which each period is the sum of events *j* within). When letting ĵbe a period of events *j*, Eq. 2 could be further rewritten as:

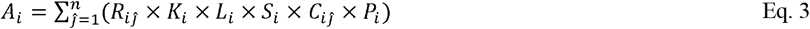

Eq. 3 forms the basis for splitting the calculation into 23 periods of 16 days, each with an individual *R*-value and an individual *C*-value (see below).

#### 2.1.3 Avoided erosion

From Eq. 1, one could calculate the effect of the vegetation and cover management in avoiding soil erosion by comparing Eq.1. with and without C and *P* factors. Where the avoided erosion in tons sediment hectare^-1^ year^-1^ (*V*_*i*_) represents the benefits of vegetation and good management practices:

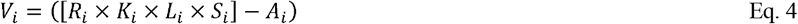

Avoided erosion is dependent on the absolute amount of possible erosion – *i.e*., if there is few potential erosion the avoided value is low, whereas potential erosion is high the avoided erosion would also be high. Therefore, this can be made into a proportion capturing the proportional effect of vegetation and cover management following:

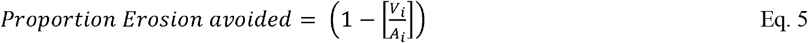

#### 2.1.4 Erosion to the streams

Subsequently, the InVEST model estimates which proportion of sediments eroded per cell would be reaching the stream, the Sediment Delivery Ratio (SDR). This is a function of the upslope area providing run-off and the downslope flow path, based among pixel connectivity. This connectivity describes the hydrological linkage between sources of sediment (from the landscape) and sinks (like streams). Higher values indicate that a greater fraction of sediment eroded from an uphill pixel is delivered to a downslope sink such as a stream (*i.e*., is more connected). Subsequently, sediment export (E) to the stream is calculated as:

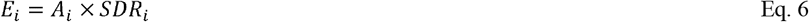

The full description and its mathematical framework are too extensive to repeat here, we point to the InVEST user guide description (NatCAP 2023).

### 2.2 InVEST Inputs not generated for this work

InVEST requires a fixed set of inputs, which are free to choose. All mapped input variables are resampled by the program to the scale of the added Digital Elevation Map. Input variables that are not specially generated for this work will be mentioned first. A second set of variables – the land cover map, the cover management factors, erosivity (*R*)– will be discussed in more length as these have been generated for this work.

#### Digital Elevation Map

We used the UK hydrological digital terrain model grids at a 50-meter resolution (Morris and Flavin 1990; 1994), this data-set is obtained under Data License agreement 1668.

#### *Soil erodibility* (K, see eq. 1)

We used the high-resolution soil erodibility map v.2014 at 500-meter resolution from ESDAC-JRC (Panagos *et al*. 2014; https://esdac.jrc.ec.europa.eu/content/soil-erodibility-k-factor-high-resolution-dataset-europe). This layer excluded 500-meters cells that were only in part land-based. Therefore, it was resampled in 25-meter gridcells without changing the value of the cells and subsequently nibbled to cover the full area of GB as set by the LCM (GitHub: Nibling&proportion.ipynb).

#### Drainages

To force the streams calculated by the DEM into observed patterns, the UK digital river centre-line network at a 1:50,000 scale was used (Moore *et al*. 1994), this data-set is obtained under Data License agreement 1668. As InVEST requires such in raster-format, these polylines were converted to 25-meter gridcells.

#### Connectivity parameters related to the SDR part of the model

1. The threshold flow value – the number of upslope pixels that must flow into a pixel before it is classified as a stream– was set at 100. This is lower than the recommended 1000 from NatCap (2023) but matches the precision in all data-sets. Sensitivity analyses showed this to have no clear influence on the validation result (less than 1% difference between 100 and 1000).
2. The Borselli *IC*_*0*_ parameter was set at 0.4671. This was estimated following 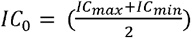 in which first as run was taken with default parameter 0.5. From the resulting grid of IC (intermediate results), the normalised maximum and minimum values were extracted and used to calculate *IC*_*0*_.
3. The Borselli k parameter was set at 2, as recommended by NatCap (2023).
4. Similarly, default recommendations were followed for the maximum SDR value that a pixel can have (set at 0.8), and the maximum allowed value of the slope length parameter (*L*) in the *LS* factor (set at 122). For the latter sensitivity analyses showed this to have no clear influence on the validation result (less than 1% difference between 122 and a maximum allowed 333 value).

### 2.3 InVEST Inputs generated for this work

#### 2.3.1 Land Cover Map Plus Crops

A comprehensive map was generated combining land cover types of Rowland *et al*. (2017; LCM 2015) with individual crop locations. Firstly, this generates the LCM input for InVEST, including relevant differences between crops via differing *C-*factor in the RUSLE equation. Secondly, this was done to allow crop specific observational assessment of these *C-*factors (see below). Here, the LCM 2015 at 25-meter raster (Rowland *et al*. 2017) was combined with Land Cover Plus Crops for 2016 (https://www.ceh.ac.uk/data/ceh-land-cover-plus-crops-2015) – the closest available full cover year to the employed LCM–. The LCM year 2015 was used to align with all other Ecosystem Services modelled under AgLand (NEC07065). A table with the crops, present in Land Cover Plus crops with their codes, can be found in Table 1.

**Table 1.**
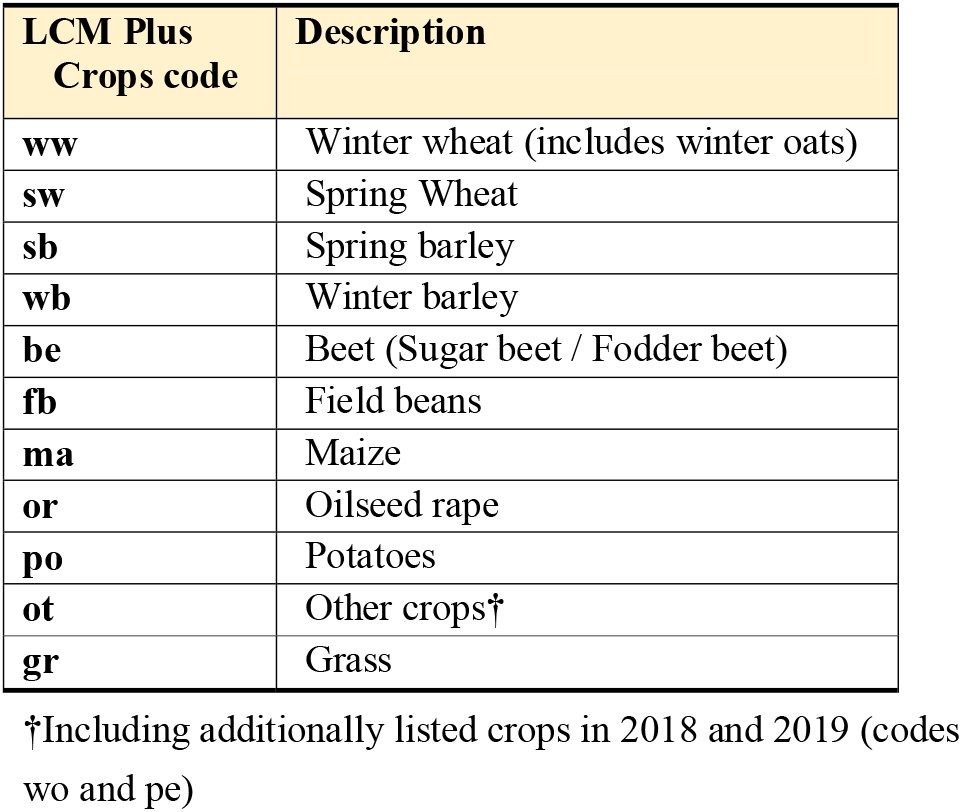
Crop descriptions and their codes in Land Cover Plus Crops.

| **LCM Plus**<br>**Crops code** | **Description** |
| --- | --- |
| **ww** | Winter wheat (includes winter oats) |
| **sw** | Spring Wheat |
| **sb** | Spring barley |
| **wb** | Winter barley |
| **be** | Beet (Sugar beet / Fodder beet) |
| **fb** | Field beans |
| **ma** | Maize |
| **or** | Oilseed rape |
| **po** | Potatoes |
| **ot** | Other crops† |
| **gr** | Grass |

†Including additionally listed crops in 2018 and 2019 (codes wo and pe)

The crop polygon layer, with LCM2007 parcel IDs (layer description), was converted to 25-meter cells and snapped to the LCM such that cells are identical and fully overlapping. Subsequently, in the combination map, the LCM categories 3 (arable) and 4 (improved grassland) were subdivided into the specific crops as present in the crop layer (see Table 1 & Table 2). The crop layer was made leading in setting a crop category or assign grassland vegetation, overriding the LCM where applicable.

**Table 2.** Employed *C-*factor for 16-day periods. (combined C and P management factors) calculated from observed NDVI (2016-2019) using van de Knijff *et al*. (2000) for the crops in the Land Cover Plus Crops and woodland categories. With added their annual mean and the values used for the baseline-run, both for the additional runs provided to detect for improvement of these observed *C-*factors and the periodicity.

| Day # | Broad' wood | Conif' wood | Winter Wheat | Summer Wheat | Winter Barley | Spring Barley | Maize | Oilseed rape | Potatoes | Field Beans | Beet | Other crops | Grass-land |
| --- | --- | --- | --- | --- | --- | --- | --- | --- | --- | --- | --- | --- | --- |
| 1 | 0.29 | 0.31 | 0.26 | 0.42 | 0.22 | 0.38 | 0.29 | 0.15 | 0.40 | 0.39 | 0.43 | 0.31 | 0.18 |
| 17 | 0.24 | 0.32 | 0.19 | 0.38 | 0.17 | 0.37 | 0.28 | 0.11 | 0.36 | 0.35 | 0.38 | 0.27 | 0.17 |
| 33 | 0.21 | 0.21 | 0.20 | 0.36 | 0.17 | 0.35 | 0.27 | 0.13 | 0.36 | 0.35 | 0.42 | 0.27 | 0.13 |
| 49 | 0.23 | 0.20 | 0.24 | 0.39 | 0.21 | 0.40 | 0.31 | 0.19 | 0.41 | 0.39 | 0.47 | 0.31 | 0.17 |
| 65 | 0.19 | 0.17 | 0.20 | 0.37 | 0.16 | 0.36 | 0.28 | 0.15 | 0.39 | 0.36 | 0.47 | 0.27 | 0.13 |
| 81 | 0.16 | 0.18 | 0.12 | 0.33 | 0.11 | 0.35 | 0.26 | 0.09 | 0.36 | 0.31 | 0.39 | 0.24 | 0.10 |
| 97 | 0.11 | 0.13 | 0.06 | 0.28 | 0.05 | 0.29 | 0.24 | 0.06 | 0.34 | 0.26 | 0.36 | 0.20 | 0.06 |
| 113 | 0.09 | 0.14 | 0.06 | 0.24 | 0.05 | 0.23 | 0.25 | 0.07 | 0.33 | 0.23 | 0.34 | 0.18 | 0.06 |
| 129 | 0.04 | 0.09 | 0.03 | 0.12 | 0.03 | 0.11 | 0.24 | 0.04 | 0.28 | 0.13 | 0.28 | 0.13 | 0.04 |
| 145 | 0.03 | 0.07 | 0.03 | 0.05 | 0.03 | 0.04 | 0.22 | 0.02 | 0.20 | 0.07 | 0.20 | 0.11 | 0.03 |
| 161 | 0.05 | 0.10 | 0.04 | 0.05 | 0.06 | 0.05 | 0.18 | 0.04 | 0.12 | 0.06 | 0.11 | 0.10 | 0.05 |
| 177 | 0.04 | 0.09 | 0.06 | 0.05 | 0.13 | 0.05 | 0.10 | 0.09 | 0.07 | 0.05 | 0.07 | 0.11 | 0.05 |
| 193 | 0.04 | 0.07 | 0.20 | 0.12 | 0.26 | 0.12 | 0.07 | 0.25 | 0.06 | 0.14 | 0.08 | 0.15 | 0.07 |
| 209 | 0.08 | 0.13 | 0.37 | 0.26 | 0.36 | 0.27 | 0.09 | 0.39 | 0.12 | 0.28 | 0.09 | 0.22 | 0.10 |
| 225 | 0.09 | 0.13 | 0.40 | 0.33 | 0.36 | 0.32 | 0.09 | 0.36 | 0.18 | 0.33 | 0.09 | 0.24 | 0.09 |
| 241 | 0.08 | 0.11 | 0.39 | 0.34 | 0.33 | 0.33 | 0.10 | 0.34 | 0.24 | 0.37 | 0.07 | 0.23 | 0.08 |
| 257 | 0.06 | 0.10 | 0.36 | 0.32 | 0.28 | 0.30 | 0.11 | 0.36 | 0.30 | 0.38 | 0.06 | 0.22 | 0.06 |
| 273 | 0.09 | 0.12 | 0.36 | 0.33 | 0.26 | 0.29 | 0.21 | 0.40 | 0.36 | 0.40 | 0.10 | 0.24 | 0.08 |
| 289 | 0.11 | 0.14 | 0.29 | 0.28 | 0.20 | 0.24 | 0.25 | 0.34 | 0.34 | 0.33 | 0.14 | 0.21 | 0.08 |
| 305 | 0.15 | 0.18 | 0.29 | 0.28 | 0.19 | 0.25 | 0.28 | 0.31 | 0.35 | 0.31 | 0.18 | 0.23 | 0.10 |
| 321 | 0.21 | 0.23 | 0.29 | 0.29 | 0.20 | 0.28 | 0.30 | 0.29 | 0.35 | 0.28 | 0.21 | 0.25 | 0.12 |
| 337 | 0.30 | 0.35 | 0.33 | 0.36 | 0.26 | 0.34 | 0.35 | 0.33 | 0.40 | 0.32 | 0.30 | 0.31 | 0.21 |
| 353 | 0.30 | 0.32 | 0.32 | 0.40 | 0.26 | 0.36 | 0.33 | 0.27 | 0.41 | 0.37 | 0.39 | 0.32 | 0.20 |
| **Annual Mean** | 0.14 | 0.17 | 0.22 | 0.28 | 0.19 | 0.27 | 0.22 | 0.21 | 0.29 | 0.28 | 0.25 | 0.22 | 0.10 |
| **Baseline-run** | 0.001† | 0.001† | 0.10‡ | 0.10‡ | 0.11‡ | 0.11‡ | 0.20‡ | 0.08‡ | 0.34‡ | 0.32† | 0.34‡ | 0.26 | 0.01† |

Following †Bakker *et al.* 2008 ‡Panagos *et al.* 2015b

Similarly, the LCM categories of calcareous, neutral, and acid grasslands were overriding by the crop layer were applicable. The codes to do so are on GitHub (CombinationLCMPlusCrops.ipynb). The combined map will be referred to as LCM+ for 2016 and is used as input into InVEST.

The same combination was made for LCM 2017 (Morton *et al*. 2020a) with Land Cover Plus Crops for 2017, LCM 2018 (Morton *et al*. 2020b) with Land Cover Plus Crops for 2018 and LCM 2019 (Morton *et al*. 2020c) with Land Cover Plus Crops for 2019. However, these additional LCM+ maps were not used as model input themselves only for multi-annual *C-*factor estimation (below). Although they could be used as model input if required. All four Land Cover Plus Crops maps were obtained under Data License agreement 1668.

#### 2.3.2 Cover management factors (C-factor)

In the baseline-run presented in October and in Panagos *et al*. (2015a; 2015b), a single value for cover management and associated tillage per crop type (*C* and *P*) was used for the full year. However, the locations for which those values are viable are less clear, often being passed over between sources (Benavidez *et al*. 2018). Here, we generated GB specific values for the cover management factor based on observed soil cover. Moreover, this project is crop oriented, hence special focus lies on such crop specific cover management factors. Importantly, crops do not cover the soil equally throughout the year. In periods there is a fuller cover whereas in *e.g*., post-harvest periods the soils could be fully exposed depending on factors like stubble management and tillage regime. In addition, the tillage support factor (*P*_*i*_) – full tillage, reduced tillage or no-tillage– is data not spatially present in GB as available data. For these reasons, we have chosen for an observational approach, assessing the combined effect of *C*_*i*_ and *P*_*i*_ factors (further referred as *C-*factor) from soil cover using satellite imagery of the Normalized Difference Vegetation Index (NDVI; van der Knijff *et al*. 2000; Benavidez *et al*. 2018). NDVI is an equation of the balance of near infrared and visible light (especially green) reflectance of the vegetation. In which unhealthy or sparse vegetation, *i.e*., no cover, reflects more visible light and less near-infrared light. Having NDVI observations throughout the year allows a periodic assessment – in 16 day intervals matching the data available from MOD13Q1 v061 (Didan 2021) – following the reasoning set by Eq. 2 and Eq. 3.

For the different crops, per 16-day period *j*, Cover management factor values were generated based on NDVI using the regression equation between NDVI and the *C-*factor following by van der Knijff *et al*. (2000):

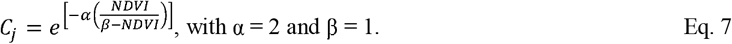

Two sets of NDVI data were used.

1. From the Sentinel satellite using Google Earth Engine extractions for 16 days intervals for the period 1-1-2016 to 31-12-2019 at a 10-meter resolution. All available maps from the Copernicus S2 data-set were selected for the given 16 days interval, with an QA60 layer cloud mask set to no-data. The NDVI value taken was the maximum value found per gridcell NDVI scores within each 16 day interval.
2. From the Terra satellite the MODIS NDVI data for the period 1-1-2016 to 31-1-2019 in 16 day intervals were downloaded at a 250-meter resolution (MOD13Q1 v061; Didan 2021; https://lpdaac.usgs.gov/products/mod13q1v061/), which have been corrected for cloud cover.

As follows from Eq. 7, the *C-*factor is inversely related to NDVI, a high *C-*value implies high erosion potential caused by lack of vegetation, whereas a low *C-*value implies low erosion capacity. *C-*factor calculations were done for all 30 combined Land Cover Plus (LCM+) classes, including 10 crop types and (semi-) permanent grassland (Table 1), for the full of Great Britain using Eq. 7. See GitHub for calculation codes (NDVI_to_Codes.ipynb).

The NDVI map was converted to *C-*factor per gridcell using Eq. 7. Subsequently, per LCM+ class, the mean value among all *C-*values was extracted using ARGIs zonal-tools with a forced 25-meter grid size environmental setting. Per period the values were averaged over the 4 years (*i.e*., the average of 4 values per period per LCM+ class). Hence, these values present a single value for GB for that land cover category (including crops) for a 16-day period as multi-year average (*C*_*ij*_ in Eq. 3). More ideally the map would be used in itself but InVEST requires these values in a biophysical table per class of the provided LCM. This calculation into *C-*factors was done for both the Sentinel and Terra data. The Sentinel data provided consistently higher *C-*factor than the Modis values (*i.e*., lower NDVI), and unrealistically high compared to *e.g*., Panagos *et al*. (2015b) as well as other values reviewed in Benavidez *et al*. (2018). Therefore, as last step, the values were averaged over both satellites per 16-day period. Table 2 provides these *C-*factors these per 16-day intervals for the 11 crops and the two woodland categories, in which the *C-*factor incorporates both *C* and *P* (technically in the Biophysical table necessary for InVEST, *P* is set to 1 and the *C-*factor is entered as *C*). Table 2 contains the first estimate of crop specific cover management factors within the UK or GB.

For all other land cover categories, either annual averages were used, or pre-set values (reviewed in Benavidez *et al*. 2018). Following Bakker *et al*. (2008), rock and urban surfaces were set to not erosible, so no sediment erosion could occur from those cells. Identically shoreline land cover (littoral and sublittoral) and water itself were set at not-erosible as no erosion into stream would occur – admittedly to the sea this could occur but that is not of focus. Annual averages were employed to avoid NDVI wrongly categorising permanent but winter brown vegetation as non-cover, such as heathlands. These other parameters are reported in Table 3. For added the annual model run (see below), *i.e*., not running the different periods but a single per year run, the per period values were averaged to get a single annual value (Table 2 – Annual mean).

**Table 3.** Employed *C-*factor (combined C and P management factors) calculated from observed NDVI. (2016-2019) using van de Knijff *et al*. (2000) for the non-crop categories as annual means values – non-periodic to avoid noise caused by non-green but fully viable vegetation. Added are the values used for the baseline-run to detect for improvement of these observed *C-*factors.

| Land cover class | C-Factor | Estimation details | Baseline-run C-factor | Estimation details |
| --- | --- | --- | --- | --- |
| Neutral grassland | 0.14 | Annual mean | 0.05 | Unmanaged grass† |
| Calcareous grassland | 0.13 | Annual mean | 0.05 | Unmanaged grass† |
| Acid grassland | 0.25 | Annual mean | 0.20 | Extensive grazing† |
| Fen, Marsh, and Swamp | 0.17 | Annual mean | 0.10 | Halfway extensive grazing and water |
| Heather | 0.28 | Annual mean | 0.20 | Extensive grazing† |
| Heather grassland | 0.28 | Annual mean | 0.20 | Extensive grazing† |
| Bog | 0.27 | Annual mean | 0.20 | Extensive grazing† |
| Inland rock | 0 | Non erodible† | 0 | Non erodible† |
| Saltwater | 0 | Water | 0 | Water |
| Freshwater | 0 | Water | 0 | Water |
| Supra-littoral rock | 0 | Shoreline only, no sediments into streams | 0 | Non erodible† |
| Supra-littoral sediment | 0 | Shoreline only, no sediments into streams | 0 | Non erodible† |
| Littoral rock | 0 | Shoreline only, no sediments into streams | 0 | Non erodible† |
| Littoral sediment | 0 | Shoreline only, no sediments into streams | 0 | Non erodible† |
| Saltmarsh | 0.22 | Annual mean | 1 | Tidal flats (Borelli *et al.* 2008) |
| Urban | 0 | Non erodible† | 0 | Non erodible† |
| Sub-Urban | 0 | Non erodible† | 0 | Non erodible† |

Following †Bakker *et al.* 2008

#### 2.3.3. Erosivity

Panagos *et al*. (2015c) provides a complete rainfall erosivity (*R* in Eq. 1) dataset for European Union, UK, and Switzerland based on REDES database with high temporal resolution rainfall measurements. The Erosivity index reflects the intensity and duration of rainfall. However, Panagos *et al*. (2015c) provides a single annual value, which does captures seasonal erosion less well (Auerwald *et al*. 2015). Following Eq.3 erosivity need to be split per period (*R*_*ij*_), such periodic values are not available from Panagos *et al*. (2015c). Therefore, we used the Gridded Estimates of Areal Rainfall (CEH-GEAR; Tanguy *et al*. 2021; https://catalogue.ceh.ac.uk/documents/dbf13dd5-90cd-457a-a986-f2f9dd97e93c) at a 1-km resolution for a 20-year period in combination with the regression approach of Ferreira & Panagopoulos (2014, overviewed in Benavidez *et al*. 2018) to convert rainfall that falls in days with more than 10mm per day to Erosivity.

This follows the following regression equation (adapted to 16 days periods from months):

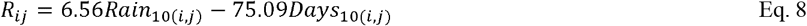

with:

*Rain*_*10(i*,*j)*_ the summed 16-day rainfall for days with ≥ 10-mm rainfall for gridcell i and interval j.

*Days*_*10(i*,*j)*_ the number of days within a 16-day period with ≥ 10-mm rainfall for gridcell I and interval j.

where:

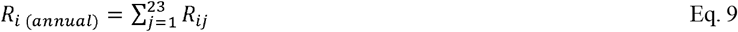

Calculations were done individually for 16-days periods for 20-years (2000-2019), subsequently per period the years were averaged. Codes are found GitHub (GEAR-ExtractionCodes.ipynb)

The advantage of this regression method is that is based on per day rainfall values (CEH-GEAR being downloadable per hour or per day), whereas following Renard *et al*. (1997) *R* is officially as sum of 30-minute erosivity sections. Data which are more scarcely present. However, the used regression parameters (6.56 and 75.09) have been trained for Portugal and were based on earlier work in Spain, with small parameter changes (Loureiro & Coutinho 2001). Portugal being a different climate than GB. It goes beyond the aim of this project to train such regression for the GB sites within the REDES data-base. In its summed form (Eq. 9), comparing to Panagos *et al*. (2015c) this generated Erosivity layer has more pronounced extremes, both on maximum (Scotland, most rain) and minimum erosivity (East Anglia, least rain). Therefore, the above approach has a bigger range compared to Panagos *et al*. (2015c). The annually summed layer is presented in Figure 1a. Whether this resembles reality can only be solved with in-field measurements.

### 2.4. InVEST Outputs used

#### 2.4.1 Main model estimates

InVEST has been run 23-times for 16-days periods, each with an individual set of *C*_*ij*_ and *R*_*ij*_ values (see above), with all other input parameters kept constant. Following Eq. 3, subsequently these 23 runs are summed into one annual figure as 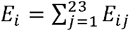and 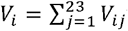with *j* the 16 days periods.

The outputs from InVEST reported here are:

*Erosion that will effectively reach the streams* in tons sediment hectare^-1^ year^-1^ per pixel estimating the amount of sediments reaching the stream and (following model assumptions) transporting these to the river outlet into the sea (*E*_*i*_, equation 6).

*Erosion avoided*, potential sediment loss that has been captured by the vegetation per pixel (*V*_*i*_). This is the difference between *A*_*i*_ calculated with and without *C*_*i*_ and *P*_*i*_ included (Eq. 4).

*The proportion avoided erosion* per pixel is independent on the absolute amount of possible erosion, and so captures proportional contribution of the standing vegetation in avoiding erosion, irrespective of the absolute amount of erosion. This proportion is generated as post-processing of InVEST output following (Eq. 5).

These outputs will be further referred to as the ‘*developed main model*’ to distinguish from the additional runs below

#### 2.4.2. Additional runs

To evaluate whether the observed crop specific *C-*factors (Table 2), the CEH-GEAR based erosivity (Figure 1a), and the periodising are actual improvements to model accuracy, four runs next to the developed main model run were generated for comparison. These will be validated along with the model developed main run (see below) with the goal of investigating the improvement generated by potential *C*_*ij*_ and *R*_*ij*_ improvements.

*The baseline-run*, the baseline set to investigate the effect of both improving on *C-*factor estimation and erosivity (*R*). This earlier model version did not include the CEH-GEAR (Tanguy *et al*.

2021) calculated erosivity but used Panagos *et al*. (2015c). As well it did not include observational *C-*factor values but using a best estimate collation from Panagos *et al*. (2015b) and Bakker *et al*. (2008). The employed *C-*factors used are listed in Table 2 (baseline) and M3. Note that since LCM+ did not fully match other parameterisations, assumptions have been made about value transfer (such as calcareous and acid grasslands being equivalent to unmanaged grass, as well as heather and bogs as extensive grazing, see Table 3).

*The full year run*. To investigate the effect of periodising, in this model version, the periods have not been used, but instead the summed annual *R*-raster (*R*_*i(annual)*_, Eq. 9) is employed. As well observed *C-*factors for crops and woodland have been averaged into full-year values, listed in Table 2 (Annual Mean).

*C changed only* (full annual only). To evaluate the improvement caused by using observational *C-* factors distinguished from the effect of a different erosivity layer compared to the baseline-run. Here in a full year run (so no periods), the CEH-GEAR estimated erosivity layer was not employed but the layer from Panagos *et al*. (2015c), as was used as input in the baseline-run.

*R changed only* (full annual only). To evaluate for the improvement caused by using a different erosivity layer distinguished from the effect of observational *C-*factors compared to the baseline-run. Here in a full year run (so no periods), CEH-GEAR estimated erosivity was added but the observational *C-*factor values were not employed but the best estimate collation that were employed input in the baseline-run.

### 2.5. Validation

The selected validation data were suspended solids (in mg/l) reported in the water quality archive from England and Wales (https://environment.data.gov.uk/water-quality/view/download) in which monitoring results are associated to GPS coordinated locations. The data-sets for 2013-2022 were downloaded and from all containing parameters the “Solids, Suspended at 105 C” category was selected for the flow type of River/Running surface water. Furthermore, only monitoring purposes were allowed removing all other goals for sampling: environmental statutory monitoring (EU, National Agency policy and UK government policy) and IPCC/IPC monitoring. Aiming at long-term averages, only water quality monitoring locations with than 4-years of data were included and with at least a total of 10 number samples. Of the 4224 unique monitoring locations, this criterium reduces the set to 320 locations. The mean concentration of suspended solids (‘sediments’) was calculated over all samples. Matlab codes are available on GitHub (WimsAverageCode.m).

Using the delineation methods described in Hooftman *et al*. (2022), polygon catchments for these 320 locations were generated (GitHub: WatershedCodes.ipynb), based on the same Digital Elevation Model as used in InVEST at a 50-meter resolution (Morris and Flavin 1990; 1994).

Subsequently, from the NFRA data-base of river discharges flow (https://nrfa.ceh.ac.uk/), the full set of 1602 locations of long-term flow discharge monitoring was downloaded. As monitoring and discharge measurement locations were only rarely performed at the exact same spot, using the ArcGIS *near* algorithm, that sediment monitoring locations that were within 1-km of a flow discharge measurement point were selected. This was done with the goal of having both solid sediment concentrations and flow discharge (reported *gdf-mean-flow* values), allowing multiplying concentrations by the amount of water flowing through the river to a total annual amount of sediments that are transported to the sea from that point (Figure 1b). It was manually checked whether the close matching locations were on the exact same river course and catchments overlap fully (except for the maximum 1-km difference), as well that no stream with an catchment of larger than 10-km^2^ was flowing into the river in-between the NFRA and water quality measurement points. This left a data-set of 178 locations with both a multi-annual mean of concentrations of suspended solids and multi-annual river discharge; those catchments do partly overlap. For every location the concentration of suspended solids (mg/l) were converted into tons of sediments flowing annually through a location by multiplying the concentration with the annual discharge in litres. The polygon delineations, that will be used for validation extractions, are the catchments generated for the sediment concentration locations in this work (Figure 1b). The NFRA delineated catchments are not used.

Following Hooftman *et al*. (2022), for the main run as well as the additional runs, predictions were obtained for each catchment in the validation dataset using the ArcGIS spatial analyst Zonal tool with a forced 10-meter grid size environmental setting to minimize edge effects, *i.e*. all predicted values were obtained by equal value resampling into 10m × 10m grid cells. Since catchments partly overlap a loop-code was used for value extractions (GitHub: WatershedCodes.ipynb & ExtractPlotValues.m).

The modelled mean value per catchment polygon was obtained and recalculated to per hectare from 25 x 25-meter values. Subsequently, both validation data (tons sediment year^-1^ in rivers) and modelled data (exported tons sediment year^-1^, summed over the area) were divided by the area of the catchment to per hectare, allowing comparison of catchments of distinct sizes at the same scale. To ensure comparability among model outputs and validation data, as well as remove any unit differences caused by scaling of input data, we standardised by normalising among the outputs prior to accuracy assessment (below). This normalisation followed Willcock *et al*. (2019) and Hooftman *et al*. (2022). To avoid impacts of extreme values without eliminating such data-points, we employed a double-sided Winsorising protocol for normalisation, using the values associated to the 2.5% and 97.5% percentiles of number of datapoints to define the 0 and 1 values (values below or above these percentiles became 0 or 1 respectively). This winsorising normalisation protocol assumes outlier data are valid, but skewed values and corrects for this by compressing the variance tails rather than trimming them (Hooftman *et al*. 2022 and citation within). This procedure and below are found on GitHub (SDRUKValidation.m).

For validation, we employed two accuracy measures (Willcock *et al*. 2019; Hoofman *et al*. 2022), which are related to different aims in modelling:

1. Comparing the rank order of predicted and validation data using Spearman ρ. Having a sufficiently trustworthy order of data-points is relevant to discover where the most important catchments are that would need erosion prevention and so prioritisation. Spearman ρ has a range from 1 (perfect positive correlation to -1 (perfect negative correlation), with the value of 0 denoting fully random. Threshold for significance for N= 178 would be around 0.30.
2. Ascertaining the absolute difference of each modelled value from its validation value using the inverse of the deviance (D↓). This is accuracy measure is relevant where modelled values are important, especially for testing for over-or under-estimations for part of modelled value ranges. This measure ranges from 0 (poor fit) to 1 (perfect fit). It has no significance statistics. For interpretation, the generally used criteria for AUC are employed, in which a result below 0.7 should be considered as likely random (Marmion *et al*. 2009; Willcock *et al*. 2019) and a value of at least 0.7 shows a “good” fit between the modelled value and the validation data.

## 3. Results and discussion

### 3.1. Description

In Figure 2, we present he first GB wide estimate of erosion and erosion retention using an (R)USLE mathematical approach using the InVEST modelling framework. Shown are the export to the stream (Figure 2a; *E*_*i*_, Methods Eq. 3) and the avoided erosion (Figure 2b & c; *V*_*i*_, Methods Eq. 4) per 50 x 50-meter pixel. Calculations included 16-day periods with each an individual set of *C-*factors and erosivity by rainfall (*R*_*ij*_ and *C*_*ij*_). The 23 periods are summed to one annual figure. Sediment export to the stream is the highest when both erosivity is high (western GB, Figure 1a) and substantial terrain differences are present (Figure 1c). Whereas in lowland eastern GB, sediment export is low. Because of that cumulative spatial effect of rainfall and terrain, differences within GB are large in predicted erosion per pixel. In Figure 3, we show two detail figures, one of Scotland with maximal erosion to the streams with peaks of predicted erosion of > 1000 tonnes per pixel per year (Figure 3a). Whereas in East Anglia exported sediments do not exceed 5 tonnes per pixel per year (Figure 3b, note the different scale).

**Figure 2.**
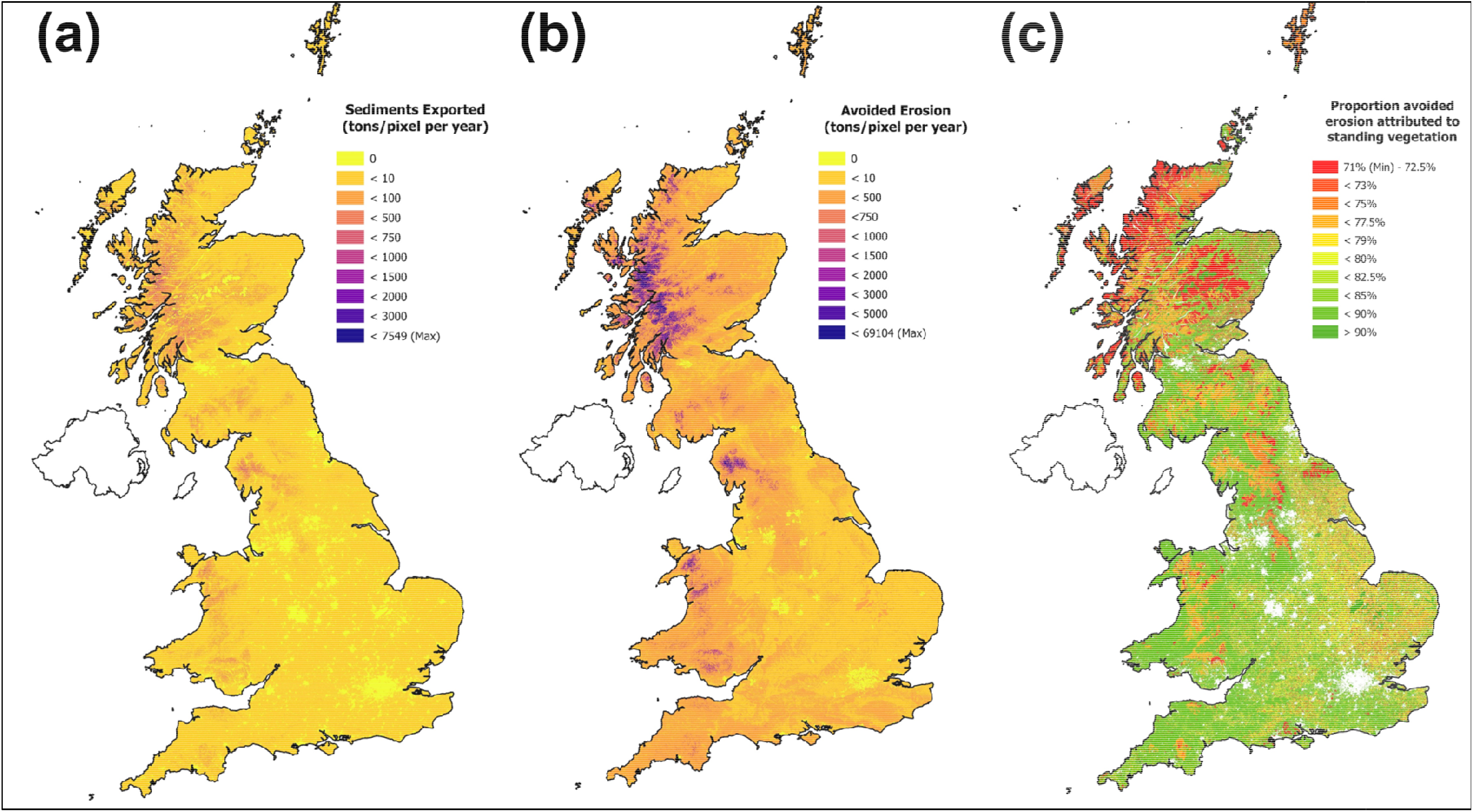
The first GB wide estimate of erosion and erosion retention using an (R)USLE mathematical approach. using the InVEST modelling framework. Shown outputs from developed model are as 50 x 50-meter pixels. (a) the estimated sediments exported per pixel to the stream (*E*_*i*_); (b) the avoided sediments exported held in place by standing vegetation (*V*_*i*_); (c) the proportion avoided sediments calculated per pixel.

**Figure 3.**
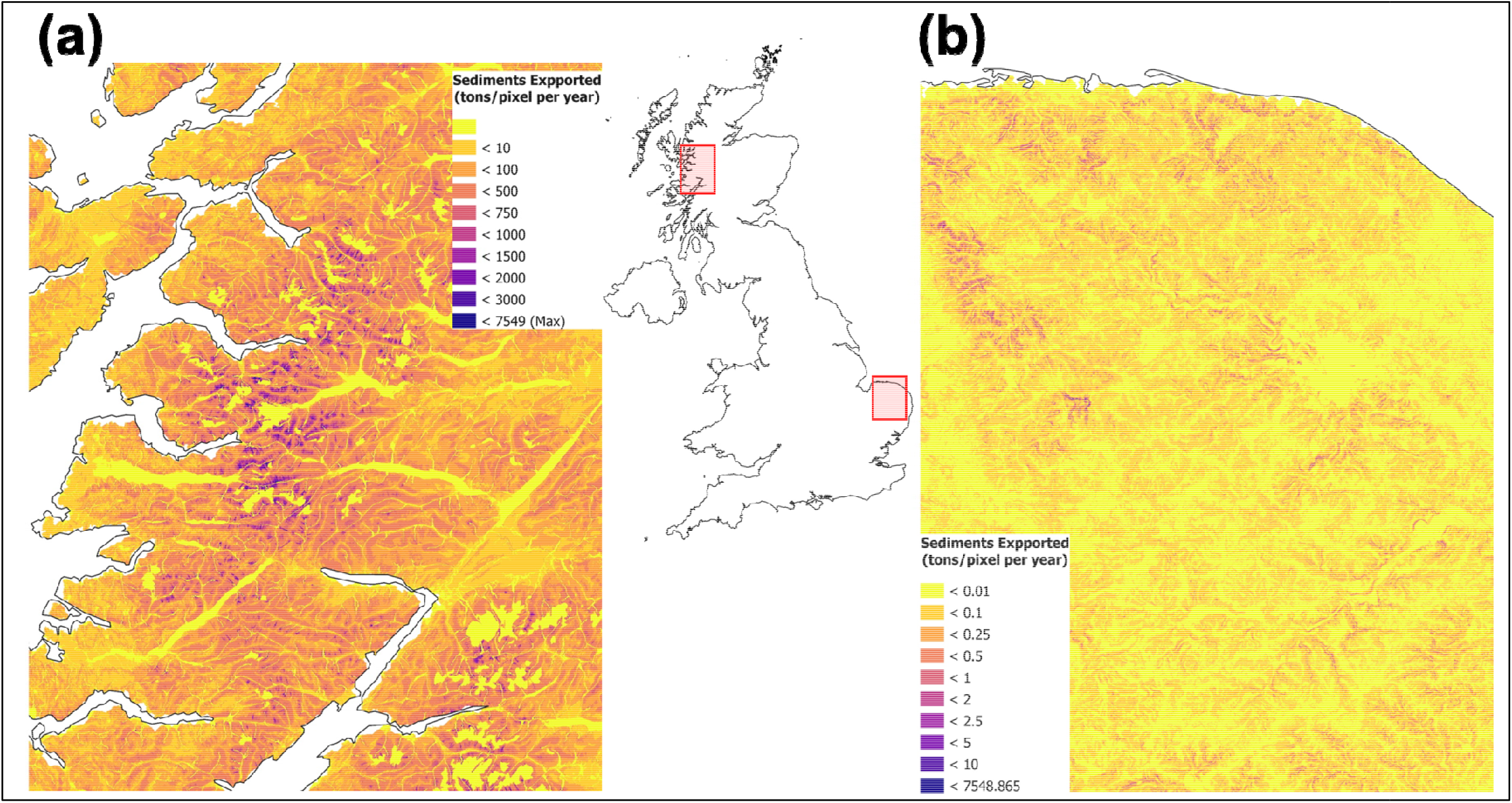
Zoomed details of the estimated sediments exported per pixel to the stream (*E*_*i*_) for (a) Scotland with high erosivity and high terrain and (b) East Anglia, being flat and with substantially less erosivity. Note the change in scales between both figures.

Avoided erosion (Figure 2b) is generally more than the occurred erosion to the stream, most potential erosion is captured by standing vegetation. Here, land cover classes with the lowest *C-*factors (Table 2 & Table 3) have the highest avoided erosion. Table 2 contains the first estimate of crop specific cover management factors within the UK or GB and is a result in itself. However, in absolute numbers avoided erosion is highly correlated to the total amount of potential erosion and therefore follows the same general spatial patterns as erosion to the streams. To make the effect of vegetation in capturing erosion more visible, Figure 2c shows the proportion erosion that is avoided, which ranges from almost 100% being avoided to just 70% avoided annually. This follows in reverse Table 2 and 3, as low *C-*factors are directly related to capture of more potential erosion by vegetation. Therefore, it is not surprising that the lowest proportion of avoided erosion is estimated in heathlands and bog areas, as *C-*factors of those land coverage categories are relatively high (see Table 3). In contrast, grass-based agriculture is highly erosion avoiding, compared to most crops. This distinction is causing the checkered patterns seen in east GB, being predominantly crop based with in between grass patches.

Similar patterning is present among the 10 crop categories themselves, *e.g*., field beans and potatoes are less good in countering in erosion (Table 3), whereas winter wheat and winter barley have a better erosion avoidance potential.

### 3.2. Validation

The developed InVEST model was validated against sediment concentrations in rivers at 178 monitoring locations, fed by their respective upstream catchments. The model significantly validated against the sediment concentrations (Table 4). The inverse of deviance (D^↓^) was with 0.73 just above the set significance threshold of 0.70 (Willcock *et al*. 2019). The Spearman ρ ranking index was highly significant (P<0.001), indicating the order of catchments is well predicted. Both indices indicate the model having a good enough predictive power. For this validation set, the most agreement is in the Lake District (high values) and in southern England (low values).

**Table 4.** Validation of the developed model for difference (D^↓^) and rank correlation against water quality monitoring data for suspended solids in rivers. Added are the validations of the additional runs answering which elements of the developed model are actual improvements.

|  | **Inverse of Deviance ($D^{\downarrow}$)** | **Spearman $\rho$ (ranking)** |
| --- | --- | --- |
| **Developed main model** | 0.73 | 0.46*** |
| **The baseline-run** | 0.74 | 0.32*** |
| **The full year run (non-periodic)** | 0.73 | 0.46*** |
| **C-changed only run (non-periodic)** | 0.76 | 0.46*** |
| **R-changed only run (non-periodic)** | 0.72 | 0.35*** |

\*\*\* $P < 0.001$

However, there is a median 115-factor difference in absolute values between observed sediments (suspended solids) detected in rivers and accumulated exported erosion following model calculations for these 178 catchments (Figure 4a and compare scales of Figure 2 and Figure 1b). This overestimation is logarithmically correlated to the amount of erosion per hectare, ranging from no to one order of magnitude overestimation at low predicted erosion to three orders of magnitude at high erosion (Figure 4a). This correlation rules out simple unit interpretation differences. Similar overestimations occur for the best available published model output, Panagos *et al*. 2015a, with a median overestimation factor of 54, which is correlated to the amount of erosion per hectare as well. Furthermore, the range of concentration of suspended solids flowing through rivers per hectare catchment is a factor 160 (maximum/minimum), whereas that is a factor 45,000 for the developed main model. It is mere speculation, how much of this severe overestimation of eroded sediments flowing through the river is caused by less-accurate model assumptions. Substantial parts are likely caused by lowered erosion detection sensitivity due to within river processes, periodicity in suspended sediment levels, and sedimentation, all potentially causing a severe lower detection of suspended sediments under high erosion levels.

**Figure 4.**
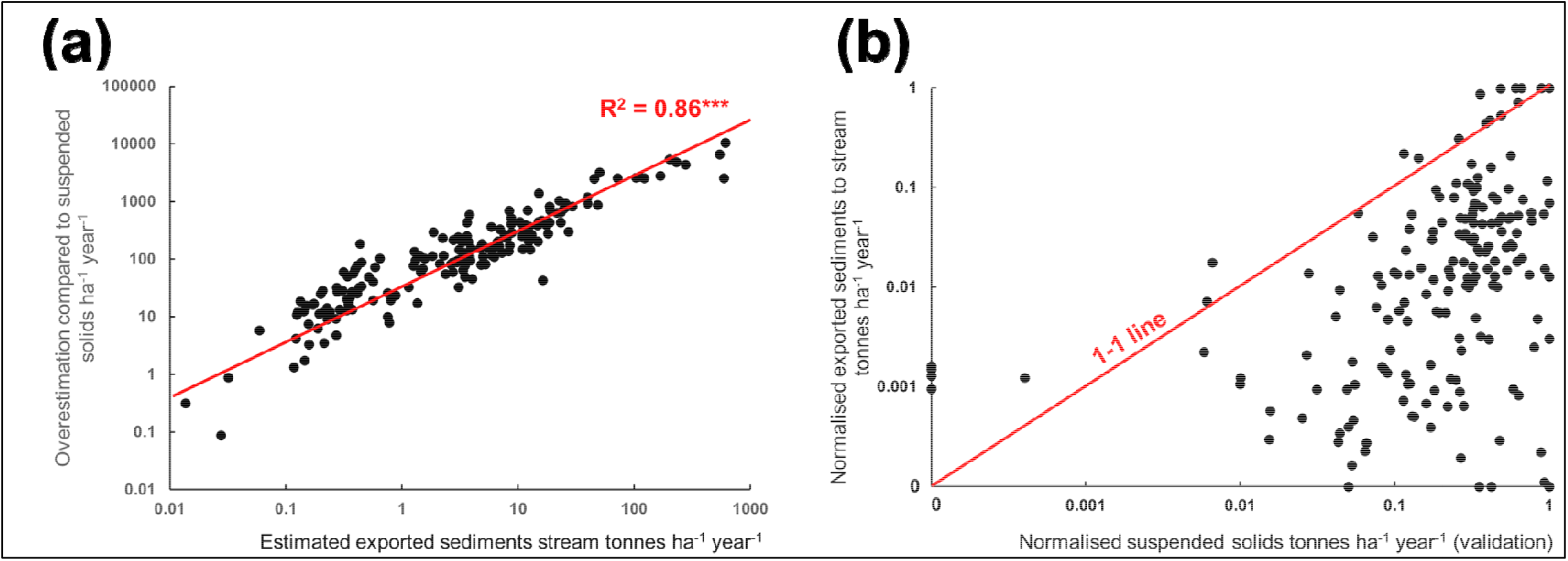
Correlations of the validation set of [suspended solids in rivers river discharge] *vs*. the estimated amount of exported sediments of the developed main model for 178 catchments, corrected for area. (a) the relationship between the overestimation, in absolute numbers, of the developed main model compared to the validation set with a highly significant log-log regression; (b) plotted normalised (winsorisation protocol) validation *vs*. estimated amount of exported sediments, with the zero deviance (1-1) Note, logarithmic scales being used to avoid showing a few high and many close zero points.

In contrast, when disregarding these differences through employing a normalised scale, for most catchments the developed model relatively underestimates the amount of erosion to the stream in tonnes per hectare catchment (Figure 4b) due to likely overestimating higher amounts of erosion skewing the distribution downwards. Compared with maximum values estimated in several catchments in the Lake District in England with higher elevation and steeper areas in, all other estimates are relatively low. This effect can be clearly seen in Figure 4b, where most data-points are substantially below the 1-1 line and most deviances are one directional. These deviances are not related to catchment size (linear regression: R^2^ < 0.01), *i.e*., the model does not consistently relatively over-or underestimates smaller catchments. Note, the highest erosion areas, in western Scotland, are not part of the validation set.

### 3.3. Improving elements

In this project three improvements were tested comparing to using the current available best estimate parameters as input in InVEST (baseline-run): C-factors estimates based on observed satellite NDVI values as proxy for soil coverages (van der Knijff *et al*. 2000; Benavidez *et al*. 2018); Erosivity (*R*) following CEH-GEAR estimates for 2000-2019 in combination with a regression approach from Ferreira & Panagopoulos (2014, overviewed in Benavidez *et al*. 2018); incorporating periodicity to allow erosion to potentially occur at moment of combined bigger erosivity and lowered soil coverage (*i.e*., especially in winter). The validation results are shown in Table 1.

Firstly, yes, the developed main model is an improvement compared to the baseline model of available best estimate parameters (Table 4: baseline-run *vs*. developed main model). The Spearman ρ ranking correlation improved from 0.33 to 0.50, *i.e*., the matching in the order of catchments from low to high sediments per hectare catchments substantially improved. Generally there are no substantial differences in the inverse of deviance (*D*^↓^) among the runs; *D*^↓^ being driven by the general underestimation (see above) rather than these improvement elements.

This improvement is fully caused by incorporating observed *C-*factors. When set as only improvement – the R layer being similar as the best-estimate baseline and no periodicity–, adding observed *C-*factor resulted in a substantial higher-ranking correlation, being the full improvement of the developed main model. This is apparent from comparing the *C-*changed only run *vs*. the baseline model in Table 4.

However, the CEH-GEAR developed Erosivity layer does not seem to perform better than the available erosivity layer from Panagos *et al*. (2015c). Rather in contrast, *D*^↓^ seems slightly lower, although the difference is small (compare *R*-changed only run *vs*. the baseline-run). Similarly, there is no difference between running the model as 23 periods of 16-days or running the model on annual averages of those periodic values (compare full year run *vs*. the developed main model). However, it could be well imagined that zooming in on specific periods, those per period estimates could be highly useful for decision making since erosion is highly variable in time, with most erosivity events (high rainfall intensity) occurring in winter in high rainfall areas in western GB.

### 3.4. Conclusion

Both the CEH-GEAR created erosivity layer as the periodic runs does not seem to be provide substantial improvement over best-estimate available data when validated against this set of suspended solids in rivers. *In contrast, the observed C-factors calculated from NDVI are a good improvement affecting the ranking order among catchments and could well be promoted as method for InVEST SDR model development and more general in RUSLE-based erosion calculations*.

## 4. Acknowledgements

This project was funded by the UKRI Landscape Decisions Agland project (NE/T000244/2) to RP. DH was subcontracted under UKCEH project number 07065. The UKCEH digital river network of Great Britain, the UK IHDTM and the UKCEH Land Cover^®^ Plus: Crops layers, were licensed to Lactuca under UKCEH Data Licence Agreement nr. 1668.

## CRediT Author Statement

**Danny A.P. Hooftman:** Conceptualisation, Methodology, Software, Validation, Resources, Data curation, Writing - Original Draft; Visualisation;

**James M. Bullock**: Conceptualisation, Supervision, Writing - Review & Editing;

**Paul M. Evans**; Conceptualisation, Project administration; **John W. Redhead:** Conceptualisation, Methodology; **Lucy E. Ridding**: Software;

**Varun Varma**: Writing - Review & Editing;

**Richard F. Pywell**: Funding acquisition, Conceptualisation, Supervision;

